# Colonization of larval zebrafish (*Danio rer*io) with adherent-invasive *Escherichia coli* prevents recovery of the intestinal mucosa from drug-induced colitis

**DOI:** 10.1101/2023.09.07.556695

**Authors:** Erika Flores, Soumita Dutta, Rachel Bosserman, Ambro van Hoof, Anne-Marie Krachler

**Affiliations:** Microbiology and Infectious Diseases Program, University of Texas MD Anderson Cancer Center UTHealth Graduate School of Biomedical Sciences, Houston, TX, USA; Department of Microbiology and Molecular Genetics, The University of Texas Health Science Center at Houston, Houston, TX, USA

## Abstract

Inflammatory bowel disease (IBD) is a broad term for a range of chronic intestinal disorders, including Crohn’s disease and ulcerative colitis. The global prevalence of IBD is rising, with over one million patients affected in the US alone. Adherent-invasive *E. coli* (AIEC) is a pathobiont frequently found in IBD biopsies. AIEC adhere to and invade epithelial cells, and can survive inside phagocytes *in vitro*. However, how AIEC contribute to IBD *in vivo* remains unclear. Here, we established a larval zevbrafish (Danio rerio) model to study the interplay between pre-existing intestinal inflammation and AIEC colonization of the gut. We used the pro-inflammatory drug dextran sulfate sodium (DSS) to induce colitis. This was followed by food-borne infection of larvae with AIEC using the protozoan *Paramecium caudatum*, a natural prey, as a vehicle.

We show that AIEC more robustly colonizes the zebrafish gut, and persists for longer, compared to non-pathogenic *E. coli*. In addition, DSS induced colitis increases both bacterial burden and persistence in the larval gut. We benchmark our model against existing rodent models using two mutants deficient in the known AIEC virulence factors FimH and IbeA, which have virulence defects in both rodent and the larval zebrafish model. Finally, we show that AIEC colonization exacerbates DSS induced colitis and prevents recovery from inflammation. In conclusion, we established a high-throughput, genetically tractable model to study AIEC–host interactions in the context of chronic inflammation.

**IMPORTANCE:** Although inflammatory bowel diseases are on the rise, a lot remains to be learned about the link between IBD severity and the underlying cause. Although host genetics, microbiome, and environmental factors have all been shown to correlate with the development of IBD, cause and effect are difficult to disentangle in this context. For example, AIEC is a known pathobiont found in IBD patients, but it remains unclear if gut inflammation during IBD facilitates colonization with AIEC, or if AIEC colonization makes the host more susceptible towards pro-inflammatory stimuli. To develop successful therapeutics, it is critical to understand the mechanisms that contribute to AIEC infections in a susceptible host. Here, we show that the larval zebrafish model recapitulates key features of AIEC infections in other animal models, and can be utilized to address these gaps in knowledge.

## INTRODUCTION

Inflammatory bowel disease (IBD) is a broad term for chronic gastrointestinal disorders, including Crohn’s disease (CD) and ulcerative colitis (UC). IBD is a major issue in industrialized nations and the number of cases in low-incidence areas is expected to keep rising (1, 2). Although the exact cause of IBD is unknown, host genetics, environmental factors, and the gut microbiota are all known disease modifiers (2).

Adherent-invasive *E. coli* (AIEC) is a bacterial pathobiont that colonizes the gut of both healthy subjects and IBD patients, but has a higher incidence in the deceased mucosae of patients with CD (21-63%) and UC (0-35.7%) (3-5). AIEC adhere to and invade intestinal epithelial cells, and survive inside macrophages without inducing host cell death *in vitro*, but how exactly they contribute to IBD is not well understood (6). It is thought that AIEC modify the pro-inflammatory environment, or inflammation facilitates AIEC colonization, because they are often isolated from lesions in patients with chronic CD as opposed to those in remission (3, 7).

Current animal models of AIEC include mice that express the human carcinoembryonic antigen-related cell adhesion molecule 6 (CEACAM6) receptor (CEABAC10 mice), conventional mice treated with broad-spectrum antibiotics, mice treated with colitis inducing agents (dextran sulfate sodium (DSS) and 2,4,6-trinitro-benzene sulfonic acid), and mice that are genetically susceptible to spontaneous colitis (8, 9). Although mice are powerful model organisms, they have some limitations that include: expensive care, long development periods, and laborious genetic manipulation. Furthermore, the scope of intravital imaging, particularly across multiple time points in mice is limited, and consequently observation of bacterial invasion, bacteria – phagocyte interactions and pathophysiological changes often require euthanasia. To address the above mentioned gaps in knowledge, we need an animal model that allows dynamic high throughput analyses and allows us to study bacteria – host cell interactions in live animals.

The larval zebrafish (*Danio rerio*) has emerged as a powerful tool to study bacterial gastrointestinal infections because the gastrointestinal tract of larval zebrafish is physiologically and functionally similar to the human intestine (10-12). Other benefits that that make zebrafish an effective high-throughput model organism include high fecundity, genetic tractability, and optical transparency throughout development into early adulthood (10). Recent studies have shown that larval zebrafish may be used to identify novel anti-inflammatory therapeutics for IBD, and that zebrafish harbor several known IBD susceptibility genes (13-15). A recent adult zebrafish model demonstrated beneficial effects of a probiotic *E. coli* strain on AIEC colonization (16).

Here, we set out to establish a model that combines a drug-inducible DSS colitis model (17) and food-borne colonization with AIEC, to investigate the interplay between host inflammation and AIEC colonization. We use the protozoan *Paramecium caudatum*, a natural prey of larval zebrafish, as a vehicle to deliver AIEC to the larval intestine, as we have previously described for other enteric pathogens (18, 19).

We benchmark this model using mutants of two AIEC virulence factors, FimH and IbeA, with known virulence deficiencies in rodent models (20, 21). We show that deletion of the type 1 pili gene (*fimH*) and the gene encoding the invasion of the brain endothelium protein A (*ibeA*) results in decreased AIEC burden, neutrophil recruitment, and epithelial damage. We also show that IbeA contributes to AIEC invasion *in vivo*. Finally, we demonstrate that colonization with AIEC hampers recovery of the intestinal epithelium from damages sustained through colitis.

## RESULTS

### Adherent-invasive *E. coli* LF82 colonizes the larval zebrafish intestine better than non-pathogenic *E. coli* MG1655

We have previously established the protozoan *P. caudatum*, a natural prey of larval zebrafish, as a vehicle for zebrafish infection with enteric pathogens and non-pathogenic *E. coli* (18, 22-24). Internalization of bacteria by *P. caudatum* and subsequent ingestion of bacteria-loaded paramecia by larvae allows for delivery of a higher bacterial dosage compared to bath immersion, which is commonly used in other zebrafish infection models including the adult zebrafish AIEC model (16, 22, 25). The uptake of bacteria-loaded paramecia by larvae is followed by digestion of the paramecia in the anterior gut and the subsequent release of bacteria into the intestine (19).

Initially, we investigated the degradation and half-life of AIEC strain LF82 following uptake into *P. caudatum* vacuoles. The uptake of AIEC by paramecia occurred rapidly, with an average burden of 339 CFUs per paramecia quantified minutes after the introduction of AIEC (**Fig.1A**). This is in accordance with other studies that show paramecia engulf their target within seconds to minutes (22, 26). The half-life τ of AIEC LF82 inside of paramecia was approximately 2.1 hours (**Fig.1A**) and was used to determine the bacterial dosage consumed by larvae following a two hour incubation with AIEC-loaded paramecia, as done previously (18). The half-life of AIEC in paramecia was similar to that reported for EHEC (22), so bacteria and *P. caudatum* concentrations were kept as described previously.

**Figure 1.**
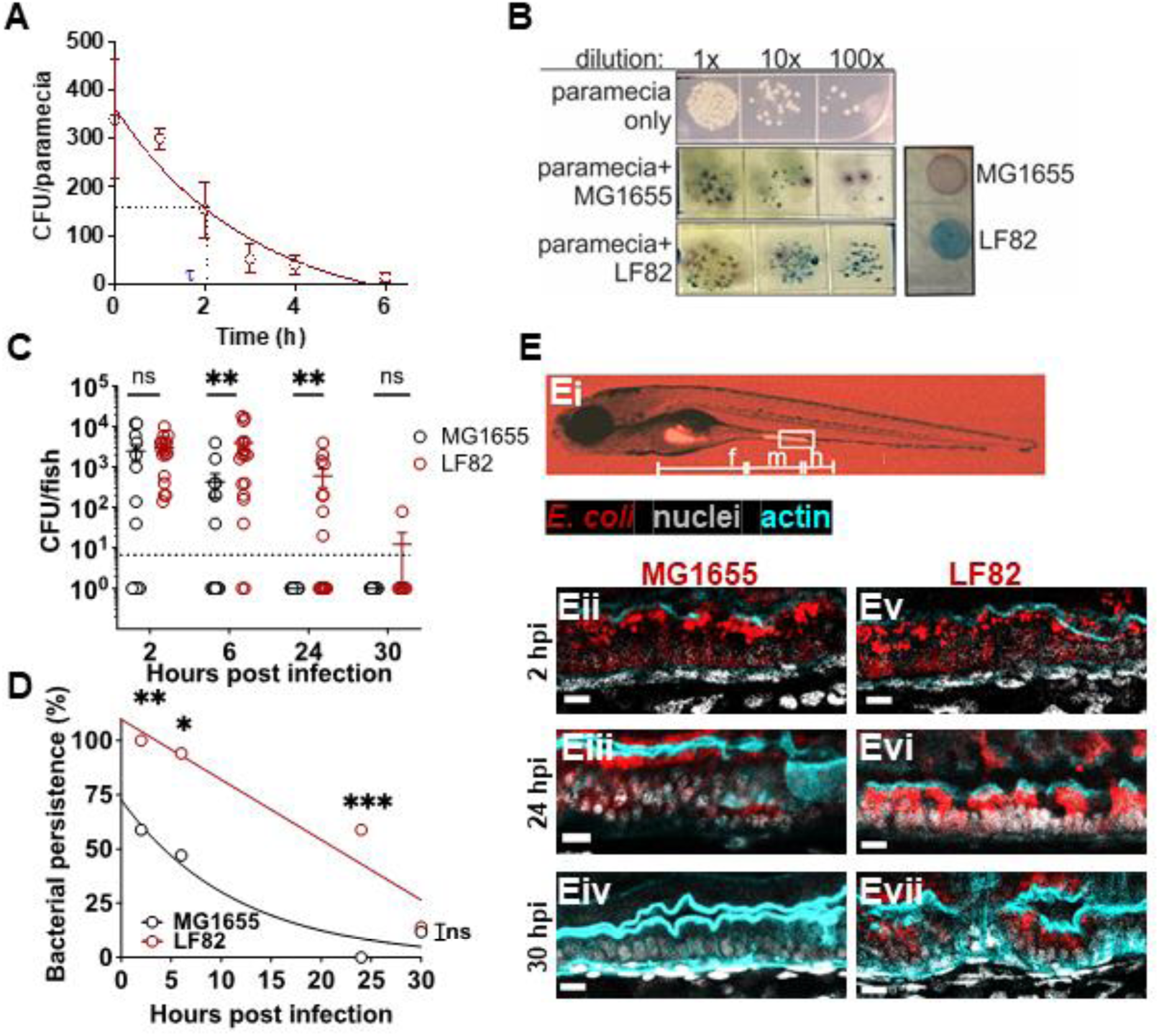
AIEC LF82 colonizes the larval zebrafish intestine better than MG1655. **(A)** AIEC-loaded paramecia sampled from 0-6 hours post incubation, and CFU/paramecia was calculated. AIEC half-life (τ) in paramecia is 2.1 hours. Data are means ± SEM, n=3. **(B)** Bacterial colonies from tissue homogenates grown on CHROMagar^TM^ O157. The zebrafish microbiota (white colonies) can be distinguished from AIEC LF82 (dark blue colonies), and *E. coli* MG1655 (mauve colonies). **(C)** Quantification of LF82 and MG1655 CFUs/fish. Fish with CFU below the detection limit (10 CFU/fish, dashed line) were annotated as 1 CFU. Data are from individual fish (n=14), and means ± SEM. **(D)** Bacterial persistence is the percentage of fish with a burden above the detection limit; n = 14. Non-linear regression, first order decay, ROUT outlier test with Q=0.2%, Paired t-test and Wilcoxon test. *, P ≤ 0.05; **, P ≤ 0.01; ***, P ≤ 0.001, ns, not significant. **(E)** Images of larvae colonized with *E. coli* (red), **(Ei)** whole larva at 10x magnification with intestinal segments (foregut (f), midgut (m), hindgut (h)) marked. Sagittal views of the midgut of larvae colonized with MG1655 **(Eii-iv)** and LF82 **(Ev-vii)** at 2, 24, and 30 hpi. Scale bars = 100 um, *E. coli* (red), phalloidin (cyan, cell outline), nuclei (DAPI, white), images are representative of n = 3;

Next, we quantified the bacterial burden of AIEC LF82 in zebrafish over 30 hpi, and used the non-pathogenic *E. coli* strain MG1655 as a control. Tissues from infected fish were homogenized and plated on CHROMagar^TM^ O157, which allowed us to distinguish AIEC LF82 (steel-blue colonies) from *E. coli* strain MG1655 (mauve), and the larva’s endogenous microbiota (white, **Fig. 1B**). Following food-borne delivery, AIEC and MG1655 were taken up by the larvae at similar concentrations (**Fig. 1C**, 2 hpi). At later time points (6-30 hpi) AIEC formed a significantly higher burden within fish than non-pathogenic *E. coli* MG1655 (**Fig. 1C**). The number of MG1655 colonized samples with a bacterial burden below the detection limit increased after 6 hpi, and by 24 hpi, no MG1655 was detected in any of the fish (**Fig. 1C)**. To get a better representation of the difference in persistence between LF82 and MG1655, the bacterial persistence was quantified as the percentage of fish that contained a burden of AIEC or MG1655 above the detection limit (≥10 CFU/fish). Although bacterial persistence decreased over time for both strains, AIEC LF82 was more persistent than non-pathogenic *E. coli* MG1655 (**Fig. 1D**). Neither colonization with MG1655 or LF82 caused any mortality throughout the experimental time course (**Fig. S1**).

We visualized the site of bacterial colonization within the zebrafish larvae using fluorescent AIEC LF82::mCherry and MG1655::mCherry strains. At 2 hpi, both strains were visible in the foregut lumen, and attached to the midgut epithelium (**Fig. 1Ei**). The localization of *E. coli* relative to the intestinal epithelium was assessed using a nuclear stain and phalloidin to outline the epithelium (**Fig. 1Eii-Evii**). High resolution fluorescence microscopy of the midgut revealed that individual AIEC and MG1655 cells localized both along the epithelial surface and inside the epithelium (**Fig. 1Eii, Ev**). By 24 hpi, luminal bacteria were no longer observed, and the burden of MG1655 had decreased (**Fig. 1Eiii**), while the LF82 burden had increased, with more invasion visible (**Fig. 1Evi**). At 30 hpi, MG1655 was no longer visible (**Fig. 1Eiv**), while AIEC LF82 was still observed within the epithelium (**Fig. 1Evii**). Taken together, these experiments showed that AIEC forms a higher burden and persists in the larval gut for longer than non-pathogenic *E. coli*, most likely by invading the intestinal epithelium.

### Larval immersion in 0.5% DSS recapitulates key morphological and pro-inflammatory features of previously described DSS colitis models

Although AIEC is found in gastrointestinal biopsies from healthy hosts, it is more prevalent in hosts experiencing chronic inflammation, such as patients suffering from IBD (27-29).

To address whether pre-existing inflammation affects the colonization and persistence of AIEC, we expanded the larval model to include drug-induced colitis. DSS is a chemical agent that induces colitis in rodent and zebrafish models. Previous studies showed that DSS causes enterocolitis in larval zebrafish, with pathologies similar to those of chronic colitis in rodents (17, 30-32).

To replicate previously described DSS colitis models, we tested different DSS concentrations and assessed larval survival, development, and inflammation (**Fig. 2, Fig. S2, and Fig. 3)**. The goal was to find a DSS dosing regimen that would induce a robust pro-inflammatory response without causing excessive mortality. Based on the experimental parameters previously described by Oehlers et al. 2012 (17), we immersed larval zebrafish in E3 media containing 0.25-0.75% DSS from 3 to 6 dpf, replacing the solution daily (**Fig. 2A).** Over the course of 4 to 10 dpf (7 days post DSS treatment), the percent survival of larvae administered 0.5% DSS decreased to 48% in comparison to untreated controls (**Fig. 2B**). The survival of DSS-treated and untreated larvae was similar at 4 and 5 dpf (1- and 2-days post treatment), however changes in the survival rate were observed at 6 dpf (3 days post treatment) (**Fig. 2B**). We observed that larval survival stabilized 3 days after the DSS was removed, and no additional mortality was observed from 7 to 10 dpf. In comparison, larvae administered 0.25% DSS had a 100% survival rate and those administered 0.75% DSS did not survive past 6 dpf (3 days post DSS exposure), (**Fig. S2A**). Consequently, we further assessed the development and inflammatory responses of larval fish treated with 0.5% DSS.

**Figure 2.**
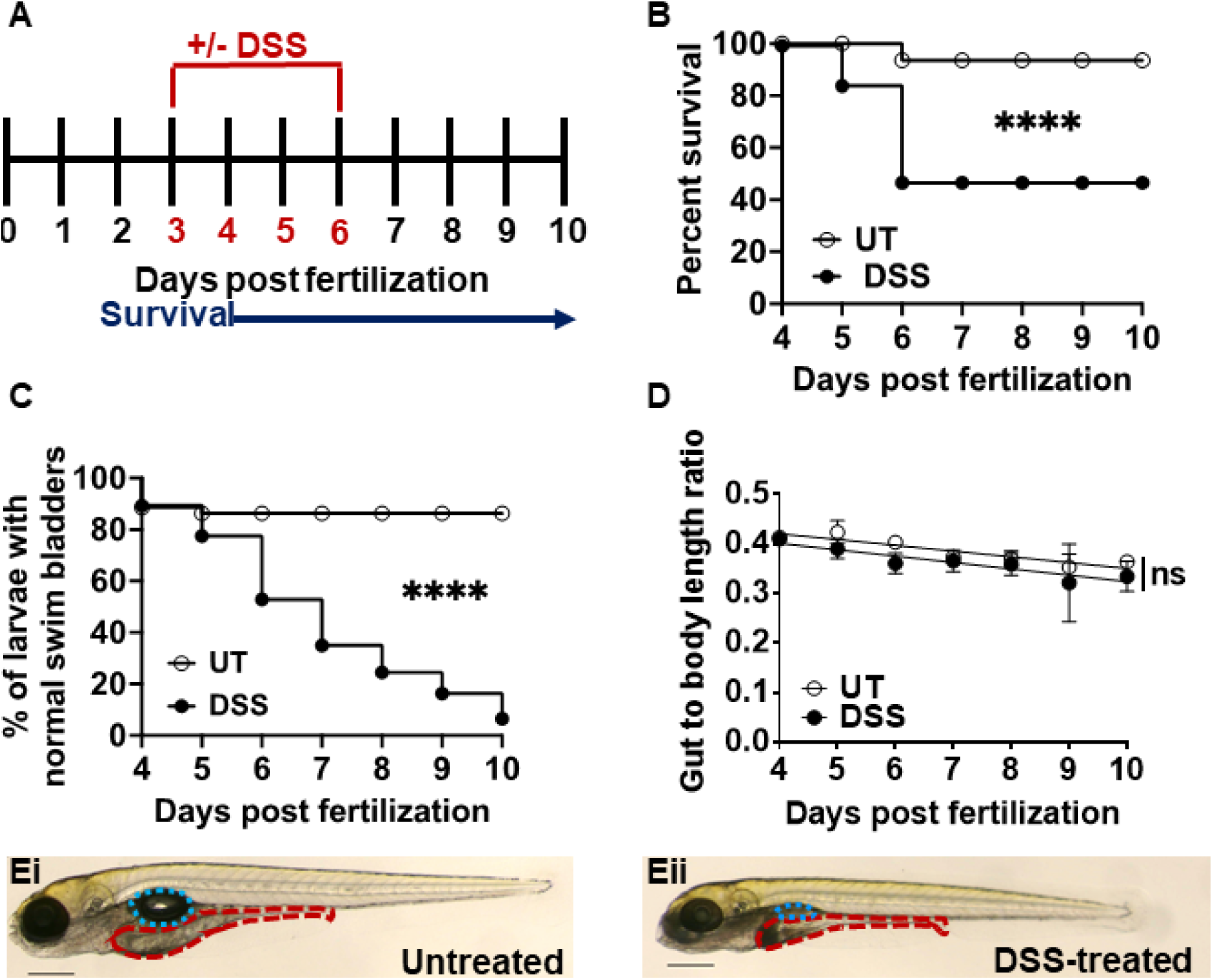
Larval zebrafish treated with 0.5% DSS have decreased survival and intestinal growth rates. **(A)** Schematic outlining timeline of DSS administration (red, 3-6 dpf) and survival experiments (blue, 1-7 days post exposure). **(B)** Survival of larvae administered 0.5% DSS (black circles) relative to untreated (UT) controls (empty circles). Data was analyzed using a Kaplan-Meier plot and Mantel-Cox test; ****= <0.0001, n=20. **(C)** Quantification of swim bladder defects in UT or DSS-treated larvae. Group differences were analyzed using Mantel-Cox test; ****, P ≤ 0.0001. n=20 **(D)** Gut to body length ratio was analyzed by linear regression; ns = not statistically significant. Data are means ± SEM from n=20; **(E)** Representative images of untreated **(Ei)** and DSS-treated **(Eii)** larvae at 6 dpf (3 days post DSS exposure), with the swim bladder (teal) and the intestine (red) outlined. Scale bar = 0.3 mm;

**Figure 3.**
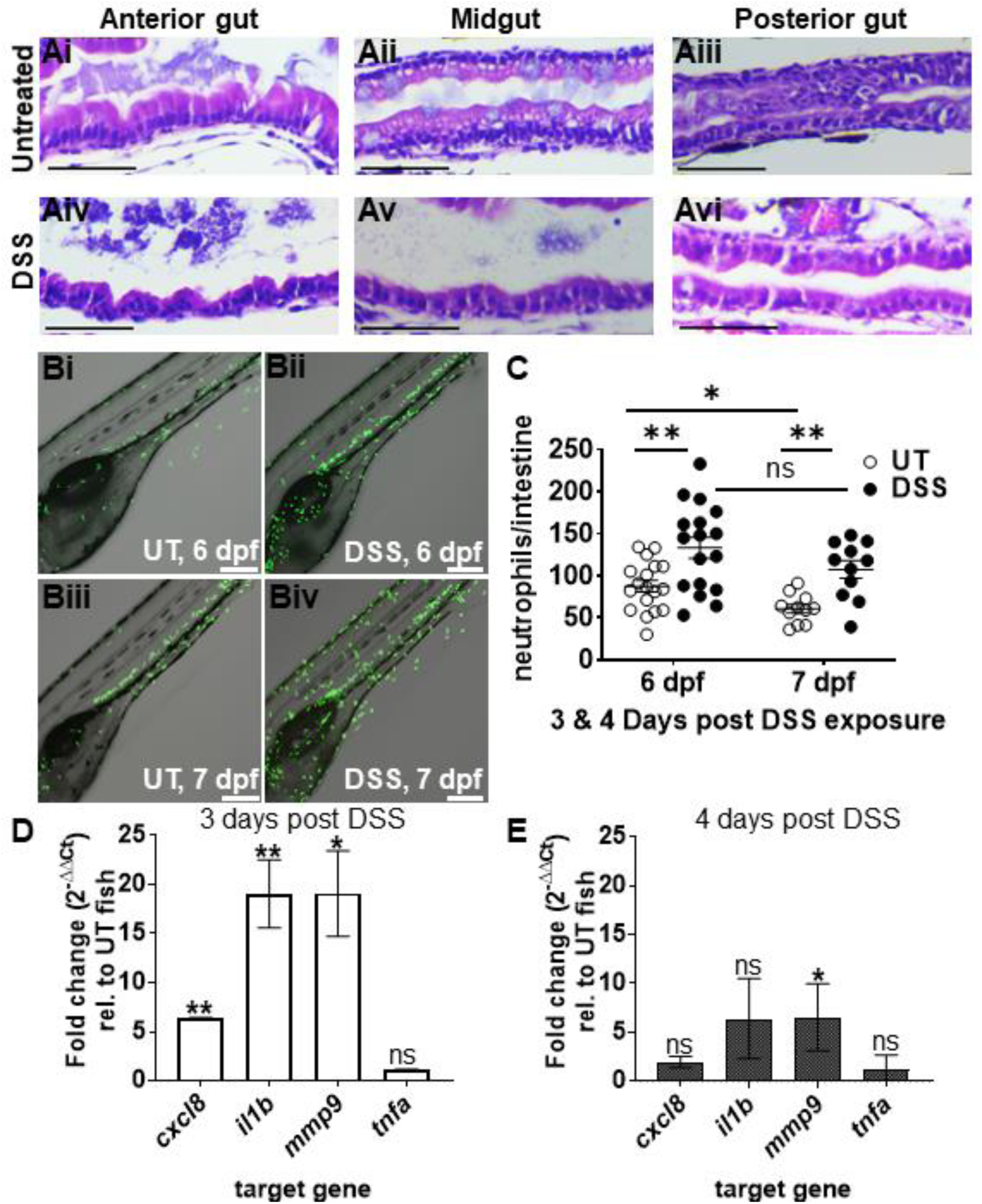
DSS causes intestinal epithelial damage and inflammation consistent with colitis. **(A)** Representative H&E stained longitudinal sections (n=4) of the anterior, mid, and posterior intestine from untreated **(Ai-iii)** and DSS-treated **(Aiv-vi)** larvae at 6 dpf; Scale bars = 50 μm. **(B)** Representative confocal images of live Tg(*mpo*::*egfp*) larvae at 6 and 7 dpf; neutrophils (green); Larvae were imaged for 18 h (3-20 hpi). Scale bars = 200 μm. **(C)** Quantification of neutrophils in the intestine at 6 and 7 dpf (3&4 days post DSS treatment); Unpaired two-tailed t-test, n ≥11, **(D)** qRT-PCR analyses of *cxcl8, il1b, mmp9,* and *tnfa* in DSS-treated larvae relative to untreated controls at 6 dpf and **(E)** 7 dpf; n=3. Unpaired two-tailed t-test. Mean ± SEM, *, P ≤ 0.05; **, P ≤ 0.01.

Prolonged treatment with 0.5% DSS led to abnormal swim bladder development over time (**Fig. 2C, E**), and slightly stunted the elongation of the larval gut and overall body length (**Fig. 2D, E and Fig. S2B, C**). Analysis of the gut to whole body ratio of untreated and DSS treated larvae suggested that DSS did not disproportionally affect gut development, but rather than shorter gut length was a consequence of overall shorter body length, since there was no significant difference in slope between untreated and DSS larvae (**Fig. 2D**). H&E staining and histology of paraffin embedded, sectioned larvae revealed normal morphology of the anterior, mid-, and posterior gut of untreated larvae (**Fig. 3A**). The intestinal epithelium was intact, with intestinal folds visible in the anterior gut, and mucus-producing goblet cells in the midgut epithelium (**Fig. 3Ai-iii**). In contrast, the epithelium was disrupted in DSS-treated larvae, with visible fraying, corrosion of intestinal folds, and epithelial detachment from the basement membrane in all three gut segments (**Fig. 3Aiv-vi**).

Next, we studied phagocyte recruitment during DSS colitis using transgenic larvae containing fluorescent neutrophils (Tg(*mpo*::*egfp*)) and macrophages (Tg(*mpeg1*::*egfp*)), respectively. Neutrophils are used as readouts of intestinal inflammation because they are the first responders to injuries and infections (33-36). Macrophages are also involved in the tissue repair and clearance of spent neutrophils, but appear at later times points (37). Live imaging of 6 to 7 dpf larvae allowed us to quantitate the number of neutrophils infiltrating the intestine. We observed that neutrophil recruitment to the intestine was significantly increased in DSS-treated versus untreated larvae at both 6 and 7 dpf (3 and 4 days of DSS treatment, respectively), (**Fig. 3B, C**). In contrast, there was no change in the number of macrophages infiltrating the gut in untreated versus DSS-treated fish (**Fig. S3**). To further evaluate pro-inflammatory signaling, we quantified the expression of the key pro-inflammatory markers interleukin 8 (*cxcl8*), interleukin-1-β (*il1b*), matrix metallopeptidase 9 (*mmp9*), and tumor necrosis factor-alpha (*tnfa*) at 6 and 7 dpf (3 and 4 days of DSS treatment, respectively). At 6 dpf, the relative expression of *cxcl8*, *il1b*, and *mmp9* was significantly increased in DSS-treated larvae compared to untreated controls, whereas *tnfa* expression remained constant (**Fig. 3D**). By 7 dpf the relative expression of *cxcl8*, *il1b* and *tnfa* was similar in DSS-treated and untreated fish, whereas *mmp9* expression remained elevated (**Fig. 3E**). Taken together, these data recapitulate key morphological and pro-inflammatory features of previously described DSS colitis models, and support our methodology of immersing larvae in 0.5% DSS from 3 to 6 dpf to induce chronic inflammation prior to introducing bacteria.

### Pre-existing DSS colitis enhances AIEC LF82 colonization, persistence, and invasion of the gut epithelium

Next, we asked whether DSS induced colitis would affect the outcome of subsequent colonization by AIEC (or the non-pathogenic MG1655 strain as a control). Following the 3-day DSS exposure, we introduced AIEC LF82 to larval zebrafish via food-borne infection (**Fig. 4A**). Larvae that had become moribund or had a deflated swim bladder following the initial DSS treatment were excluded from subsequent infection experiments. At 2 hpi, the AIEC burden in DSS colitis fish was similar to the AIEC burden in untreated fish (**Fig. 4B**). However, the burden of AIEC in DSS-treated larvae was higher than that of the untreated controls at 6 and 12 to 48 hpi (**Fig. 4B**). Further, the persistence of LF82 in DSS-treated larvae was significantly higher compared to untreated fish (**Fig. 4C**). Together, these data suggest that pre-existing inflammation enhances the burden and persistence of LF82 in the intestine of larval zebrafish. These results are also in accordance with those of published murine studies that show that AIEC persists longer in mice with IBD compared to healthy controls (38-40).

**Figure 4.**
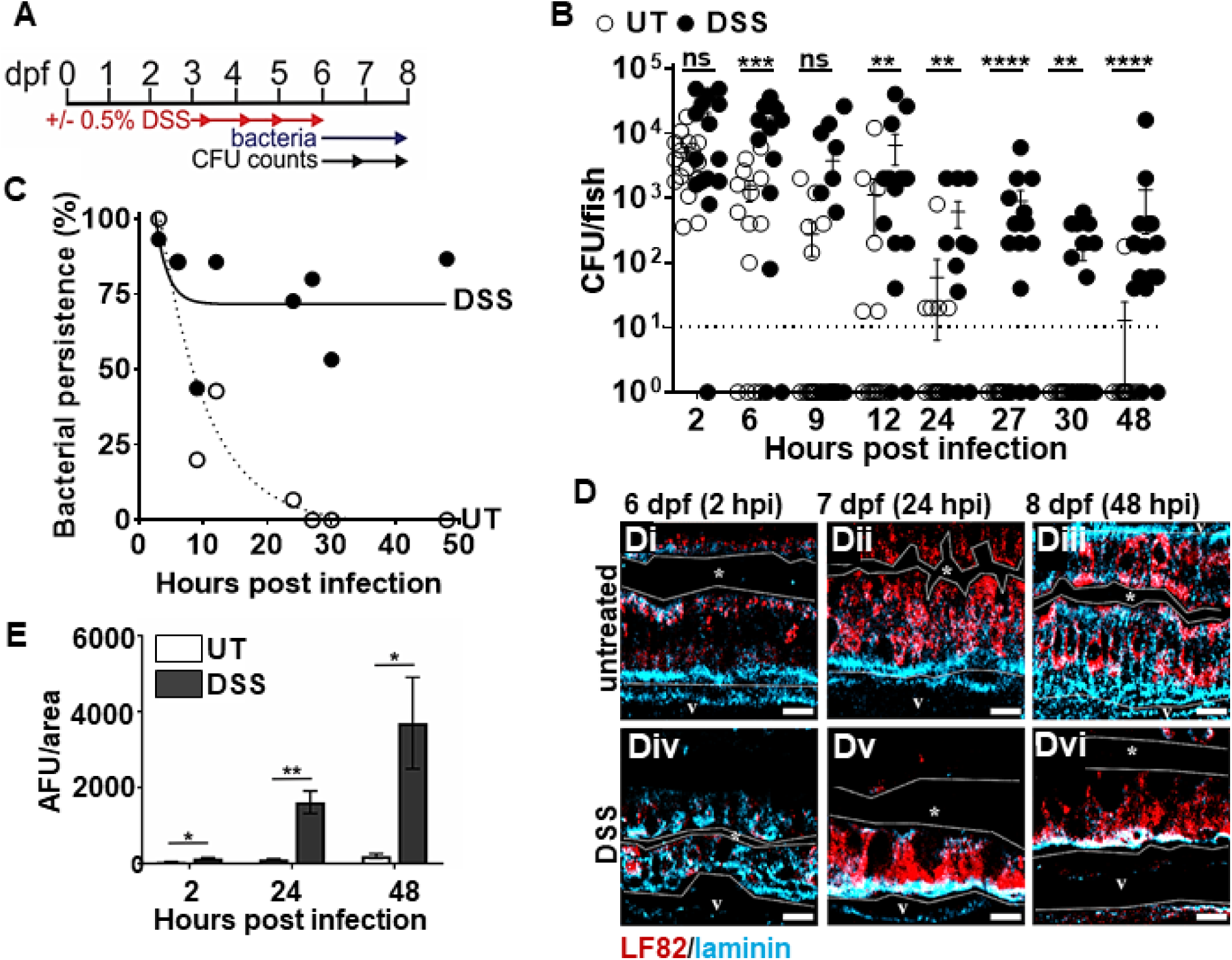
Pre-existing inflammation enhances the colonization, persistence, and invasion of AIEC LF82. **(A)**Timeline of DSS administration, infection of larvae, and sampling for CFU counts. **(B)** Quantification of LF82 CFUs per larvae with and without DSS treatment, n≥ 17; fish with CFU below the detection limit (10 CFU/fish, dashed line) were annotated as 1 CFU. **(C)** Bacterial persistence (% fish with a burden of AIEC above the detection limit); Non-linear regression first order decay, ROUT outlier test with Q=0.2%. (**D**) LF82 (red) in the mid-intestine of UT **(Di-iii)** and DSS-treated **(Div-vi)** larvae relative to the basement membrane (blue) from 2 to 48 hpi or 6-8 dpf. The dotted white line outlines the intestinal epithelium and separates it from the lumen, indicated by *, and the blood vessel below the basement membrane (v). Scale bars = 10 μm**. (E)** Quantification of red fluorescence intensity (AFU) (representing AIEC) in the vasculature (v) at 2, 24, and 48 hpi, n= 6; *, P ≤ 0.05; **, P ≤ 0.01; ***, P ≤ 0.001; ****, P ≤ 0.0001; ns, not statistically significant.

To investigate whether pre-existing inflammation enhances bacterial colonization in general, or specifically for AIEC, the colonization patterns of MG1655 in DSS-treated larvae were also assessed. The burden and persistence of LF82 were significantly higher than those of MG1655 in DSS-treated fish at 2, 6, 24 and 48 hpi (**Fig. S4A, B**). These results demonstrate that pre-existing colitis enhances the burden of both AIEC and non-pathogenic *E. coli*, and that AIEC LF82 still colonized and persisted better compared to non-pathogenic *E. coli* in fish with colitis.

Colitis damages the mucosal barrier and enhances intestinal permeability, allowing for increased bacterial invasion (17, 41, 42). Therefore, we asked whether pre-existing colitis would affect AIEC invasion in our model. DSS-treated and untreated larvae were infected with LF82, euthanized at 2, 24, and 48 hpi, and laminin and DAPI stained to assess the localization of LF82::mCherry relative to the intestinal lumen, epithelium, and underlying vasculature (**Fig. 4D**). At 2 hpi, LF82 cells were present within the epithelium of untreated and DSS-treated zebrafish, and had begun to invade the underlying vasculature in DSS-treated but not in control fish (**Fig. 4Di, 4Div, Fig. 4E**). At 24 hpi, individual bacterial cells remained visible in untreated larvae, whereas large bacterial aggregates were observed within the epithelium of DSS-treated fish (**Fig. 4Dii, 4Dv**), and increased bacterial invasion of the underlying vasculature was measured in DSS fish, but not untreated controls (**Fig. 4E**). By 48 hpi the AIEC burden within the epithelium had lowered (**Fig. 4Diii, 4Dvi**), but invasion of the vasculature in DSS treated fish was further elevated (**Fig. 4E**). Together, these data suggest that pre-existing colitis facilitates bacterial colonization and persistence, and exacerbates invasion of the bloodstream by AIEC.

### AIEC LF82 exacerbates intestinal inflammation in DSS-treated larvae

Murine studies show that colonization of AIEC LF82 exacerbates intestinal inflammation in DSS-treated animals and causes an immunopathology similar to that observed in IBD patients (40, 43, 44). Thus, we investigated whether AIEC could exacerbate inflammation in larvae with pre-existing DSS colitis. Untreated and DSS-treated larvae fed the paramecia vehicle only (uninfected) were used as controls and compared to AIEC-infected fish **(Fig. 5)**. The midgut of untreated fish colonized with LF82 contained an increased number of mucus secreting goblet cells at 2, 24, and 48 hpi compared to control fish (**Fig. 5A** vs **B**, cells containing clear/light blue mucus droplets) (45). Increased goblet cells were also observed in the posterior gut of untreated larvae infected with LF82 from 2 to 48 hpi (**Fig. S5F**).

**Figure 5.**
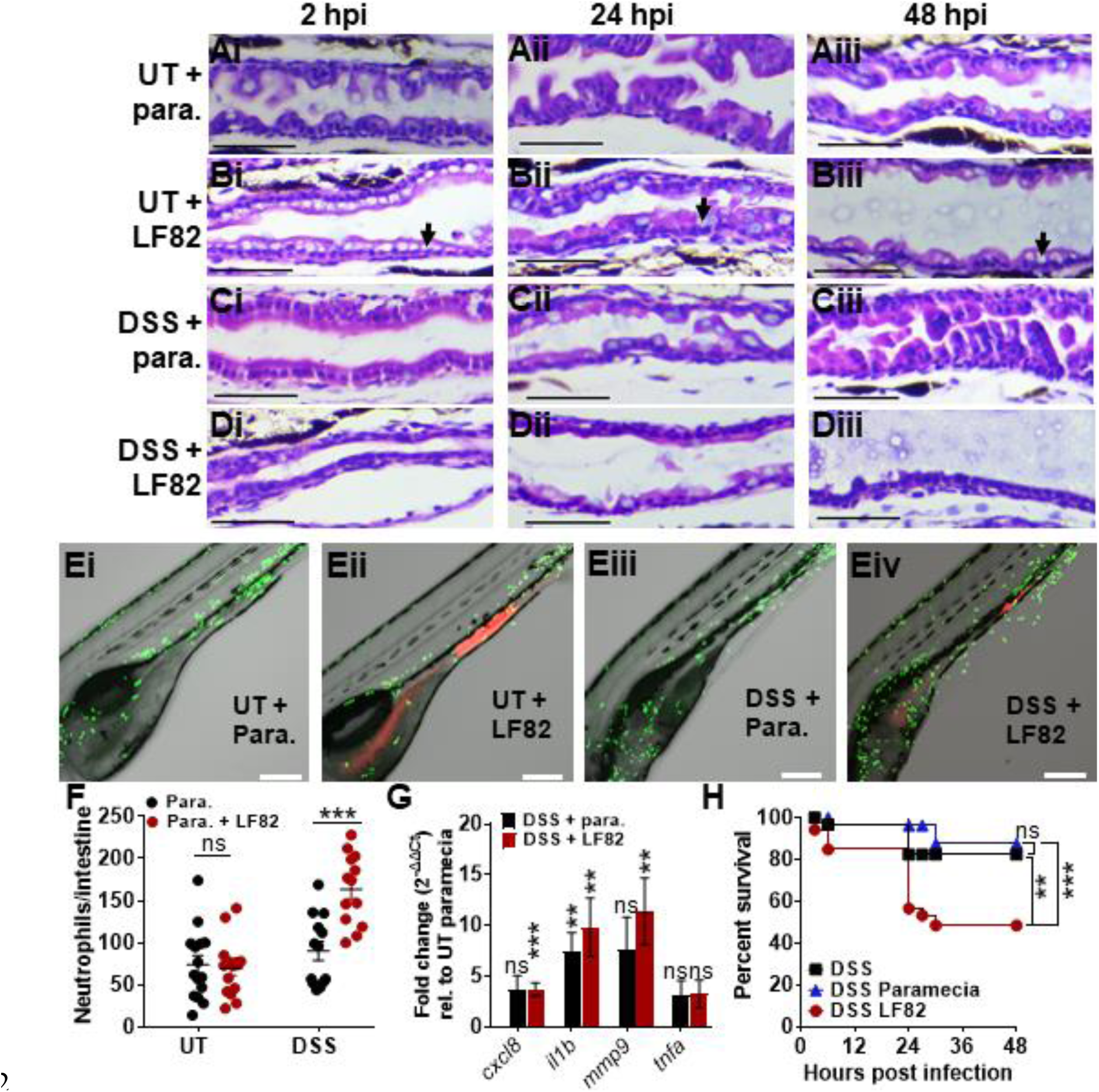
AIEC LF82 exacerbates intestinal inflammation in DSS-treated larvae. **(A-D)** H&E-stained longitudinal sections of the mid-intestine of larvae without **(A, B)** and with **(C, D)** prior DSS treatment, fed paramecia (para.) alone **(A, C)** or paramecia containing LF82 **(B, D)** at 2 **(i)**, 24 **(ii)**, and 48 **(iii)** hpi, n= 3. Black arrows point to goblet cells. Scale bars = 50 μm. **(E)** Representative confocal images of Tg(*mpo*::*egfp*) larvae fed paramecia only **(i, iii)** or LF82 **(ii, iv)** at 6 dpf. Larvae were imaged for 18 h (3-20 hpi), neutrophils (green) and bacteria (red). Scale bar = 200 μm. **(F)** Quantification of neutrophils per intestine in UT- and DSS-treated fish fed with paramecia only (black) or para. containing LF82 (red); n ≥10. **(G)** qRT-PCR analyses of *cxcl8, il1b, mmp9,* and *tnfa* in DSS-treated (red) larvae infected with LF82 and DSS-treated larvae fed paramecia (black) relative to UT paramecia controls (1-fold) at 6 dpf, n=7. Unpaired two-tailed t-test. Mean ± SEM. **(H)** Survival of DSS-treated larvae that were uninfected (black), fed paramecia control (blue), or para. containing AIEC (red). N = 17. Kaplan-Meier and Mantel-Cox test, followed by a Bonferroni correction test. **, P ≤ 0.01; ***, P ≤ 0.001; ns, not statically significant.

Following DSS treatment, we observed corrosion of intestinal folds in the midgut ((**Fig. 5Ci**) and anterior gut (**Fig. S5C**) at 6 dpf (3 days post-DSS treatment). In uninfected fish, these folds were partially restored at 7 and 8 dpf (1-2 days after DSS treatment had stopped, **Fig. 5Cii-iii, Fig. S5C**), suggesting that the damaged intestinal epithelium can recover from colitis.

In contrast, DSS-treated larvae infected with LF82 were unable to fully recover from colitis by 48 hpi, since the anterior and midgut did not recover the original intestinal fold architecture and exhibited a thinner epithelial cell layer compared to DSS-treated larvae that were not infected (**Fig. 5D vs C**, **Fig. S5D vs C**). LF82 colonization did not disrupt intestinal folds in the absence of DSS colitis (**Fig. 5B and Fig. S5B**). Together, these data suggest that AIEC LF82 alters the architecture of the intestine of larvae, in untreated fish LF82 increases goblet cell number, and in DSS-exposed fish it prevents epithelial healing. The increased presence of mucin-producing goblet cells may indicate a host-defense response to fight off bacterial infections whereas flattening of the intestinal villi may be due to inflammation (46).

To further examine the effect of LF82 on inflammation, neutrophil recruitment was assessed, and induction of inflammatory markers was quantified using qRT-PCR. In untreated fish, AIEC colonization did not affect neutrophil recruitment to the gut (**Fig. 5E, F**). Similarly, in uninfected fish, neutrophil recruitment to the intestine was unchanged following DSS treatment (**Fig. 5E, F**). In contrast, DSS treatment and subsequent AIEC colonization had an additive effect and increased neutrophil recruitment (**Fig. 5E, F**). Macrophage recruitment to the intestine was not significantly affected either by DSS treatment or AIEC (**Fig. S6**). Expression of inflammatory markers *cxcl8, il1b*, and *mmp9* was slightly elevated following DSS treatment alone, and significantly increased in DSS colitis fish colonized with AIEC (**Fig. 5G**). Comparison of marker expression following AIEC colonization of untreated or DSS colitis fish further showed that DSS colitis and AIEC infection have an additive effect on pro-inflammatory signaling (**Fig. S7**).

The observed increase in epithelial damage and pro-inflammatory response following LF82 infection in DSS colitis fish may contribute to the increase in mortality of DSS-treated larvae infected with AIEC LF82, relative to DSS alone or DSS larvae fed paramecia only (**Fig. 5H**). Together these data suggest that AIEC colonization in healthy fish causes little epithelial damage and inflammation, but exacerbates inflammation and tissue damage in hosts with pre-existing colitis.

### FimH and IbeA contribute to AIEC virulence in larval zebrafish

Next, we investigated whether the larval zebrafish model is suitable for the characterization and/or identification of virulence factors involved in *in vivo* infections by characterizing the phenotypes of two known AIEC virulence factors, FimH and IbeA, as a benchmark. FimH is the terminal subunit of type I pili and binds collagen type I and type IV, laminin, fibronectin, and mannosylated glycoproteins (47). FimH of AIEC LF82 adheres to the human CEACAM6 receptor that is abnormally expressed in the ileum of CD patients and expressed in transgenic CEABAC10 mice (8, 21). It is hypothesized that the presence of CEACAM6 receptors in a host promotes the colonization of AIEC and indirectly contributes to intestinal inflammation, since binding of AIEC to CEACAM6 through FimH triggers intestinal inflammation in CEABAC10 mice (48). IbeA is an invasin and outer membrane protein conserved in the *E. coli* phylogenetic group B2, which includes avian pathogenic *E. coli*, newborn meningitis-causing *E. coli*, and AIEC strains NRG857C and LF82 (20). BLAST analyses show that the IbeA protein in these pathogenic *E. coli* strains are 100% identical (data not shown). IbeA binds to vimentin found in macrophages, fibroblasts, and endothelial cells, and mediates the invasion of Caco-2 and M-like cells by AIEC strain NRG857c (20).

To investigate whether FimH and IbeA play a role in the colonization and invasion of AIEC LF82 in zebrafish larvae, these genes were deleted from the parent strain and complemented by inserting *fimH* or *ibeA* with their endogenous promoters into the chromosome. Deletion and complementation of either gene did not affect the overall growth of AIEC LF82 (**Fig. S8**). There were no fortuitous mutations identified in the deletion and complement strains, which were subjected to whole genome sequencing.

Deletion of *fimH* but not of *ibeA* significantly increased larval survival, and the defect was restored in the LF82Δ*fimH:fimH* complementation strain (**Fig. 6A, B**). *FimH* and *ibeA* deletion and complementation strains were taken up into the larval gut at similar levels than the wild type strain (**Fig. 6C**, **D**, 0 hpi). Interestingly, deletion of either *fimH* or *ibeA* initially increased AIEC colonization, but led to a colonization defect at 48 hpi. Complementation of *fimH* and *ibeA* restored wild type colonization levels (**Fig. 6C, D**). Bacterial persistence was unaffected by *fimH* deletion (**Fig. 6E**), but decreased upon deletion of *ibeA* (**Fig. 6F**). Next, we asked whether the deletion of *fimH* or *ibeA* affected the invasion of the epithelium by AIEC. Infected larvae were euthanized, fixed, and stained with anti-laminin and DAPI to visualize the localization of LF82Δ*fimH:mcherry* and LF82Δ*ibeA:mcherry* and complementation strains over the course of 48 hpi. Deletion of either *fimH* or *ibeA* caused a transient increase in bacterial burden at 2 hpi (**Fig. 7A-Ei**), followed by significantly decreased colonization at 24-48 hpi, (**Fig. 7A-Eii and iii**) consistent with the CFU burden data (**Fig. 6C, D**). Interestingly, while the *fimH* mutant was still able to invade the epithelium, the *ibeA* mutant mainly colonized and formed aggregates at the epithelial surface (**Fig. 7D**). Complementation of *fimH* and *ibeA* restored wild type adherence and invasion (**Fig. 7C, E**). These data suggest that FimH and IbeA both contribute to aspects of pathogenesis, but play distinct roles in bacterial adherence and invasion.

**Figure 6.**
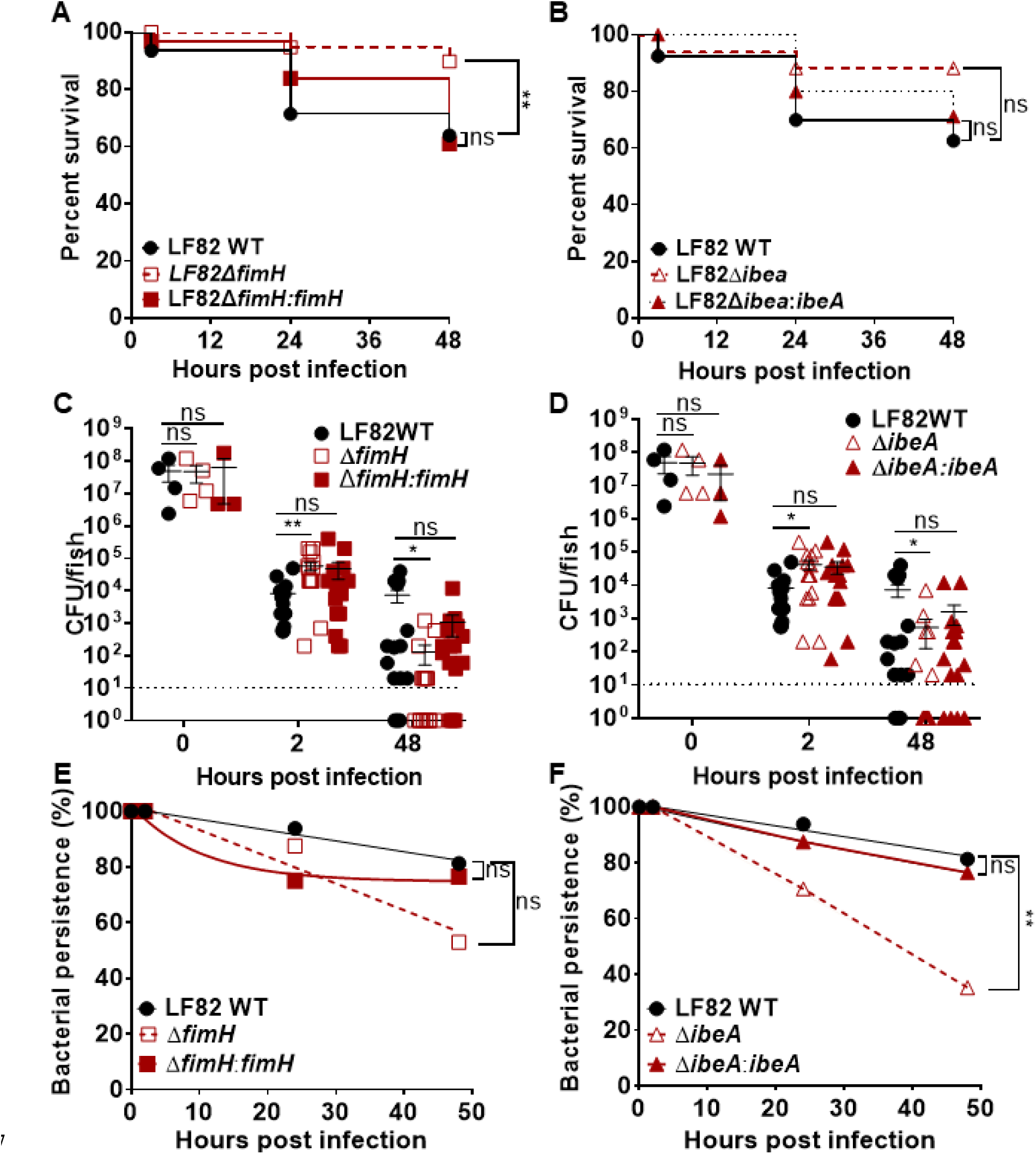
Effects of *fimH* and *ibeA* deletion on survival, burden, and AIEC persistence in larvae. Survival of larvae infected with **(A)** LF82 wild-type (WT), LF82Δ*fimH*, LF82Δ*fimH:fimH* or **(B)** LF82, LF82Δ*ibeA*, LF82Δ*ibeA:ibeA* at 2, 24, and 48 hpi. Kaplan-Meier and Mantel-Cox test, followed by a Bonferroni correction test, n=20. Quantification of bacterial burden and persistence of **(C, E)** LF82, LF82Δ*fimH*, LF82Δ*fimH:fimH*, or **(D, F)** LF82Δ*ibeA*, and LF82Δ*ibeA:ibeA* in DSS-treated larvae from 2-48 hpi. Fish with CFU below the detection limit (10 CFU/fish, dashed line) were annotated as 1 CFU. Significance of difference in burden was analyzed using a Kruskal-Wallis test, n ≥ 16. Bacterial persistence (percent of fish with a burden of AIEC above the detection limit) was analyzed using a log-rank test. Non-linear regression, first order decay graph used to model bacterial persistence. *, P ≤ 0.05; **, P ≤ 0.01; ns, not significant;

**Figure 7.**
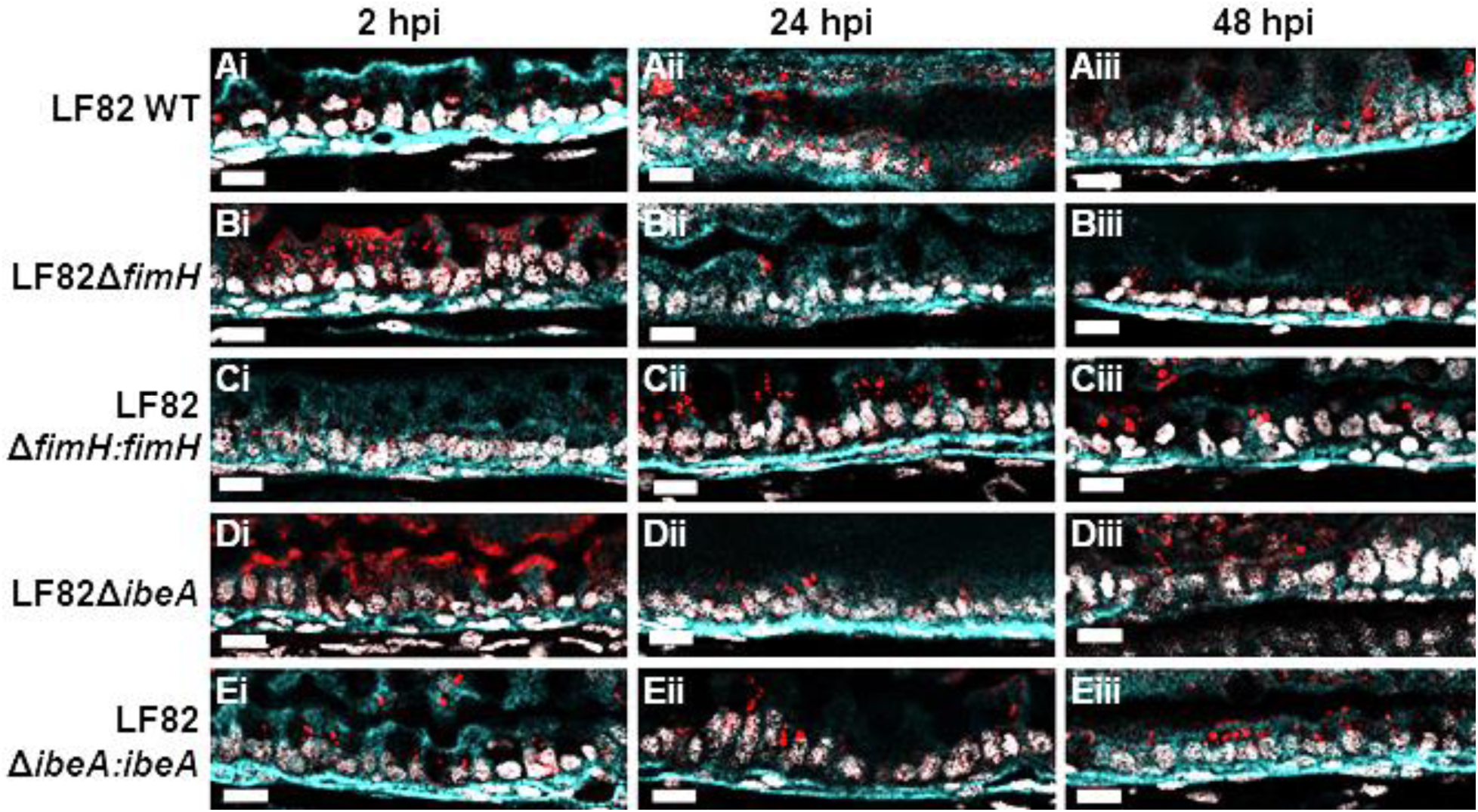
Deletion of *ibeA*, but not *fimH*, results in aggregation and retention of AIEC LF82 on the epithelial surface. Representative sections (n=3) of the mid-intestine of larvae infected with **(A)** LF82 WT, **(B)** *LF82ΔfimH*, **(C)** *LF82ΔfimH*:*fimH*, **(D)** LF82Δi*beA:ibeA*, or **(E)** LF82Δi*beA:ibeA* at **(i)** 2, **(ii)** 24, and **(iii)** 48 hpi. AIEC LF82 (red), laminin (cell surface, cyan); nuclei (DAPI, white). Scale bars represent 10 μm.

### FimH and IbeA contribute to pro-inflammatory response during AIEC colonization and prevent epithelial recovery from colitis

Since both FimH and IbeA are bacterial surface proteins, we next asked if they contribute to the pro-inflammatory response to AIEC colonization in DSS colitis fish. Histology of midgut sections from infected DSS colitis fish showed that colonization with wild type or complementation strains prevented recovery from DSS colitis, and corrosion of intestinal folds persisted even 2 days after DSS treatment had been discontinued (**Fig. 8A**, 24-48 hpi). In contrast, healthy epithelial morphology was restored following infection with either *fimH* or *ibeA* deletion strains (**Fig. 8B, D**). Lastly, we studied how FimH and IbeA contribute to AIEC immunogenicity, by quantifying neutrophil recruitment to the gut. Fish infected with LF82 WT recruited more neutrophils to the intestine compared to either uninfected, paramecia-fed fish, or fish harboring LF82Δ*fimH* and LF82Δ*ibeA* (**Fig. 8F**). Complementation of *fimH* resulted in increased neutrophil recruitment similar to or in the case of ibeA, more neutrophil recruitment than wild type infection. Taken together, these data suggest that both FimH and IbeA contribute to pro-inflammatory signaling in response to AIEC infection, and contribute to attenuation of epithelial recovery in DSS colitis fish.

**Figure 8.**
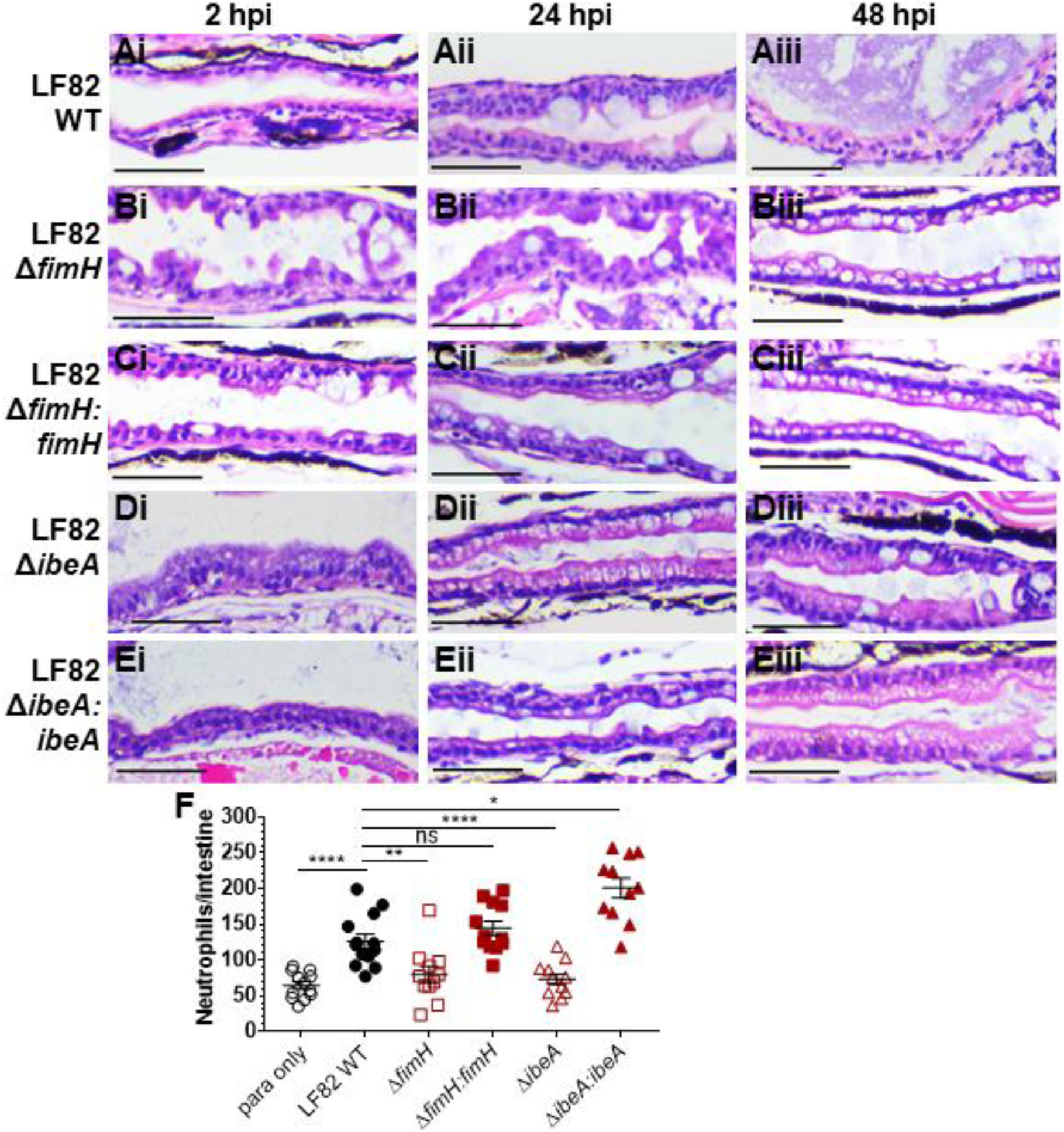
Deletion of *fimH* and *ibeA* in AIEC LF82 results in decreased tissue damage and neutrophil recruitment to the intestine compared to LF82. H&E longitudinal sections of the mid-intestine of larvae infected with **(A)** LF82 WT, **(B)** LF82Δ*fimH,* (C) LF82Δ*fimH:fimH,* (D) LF82Δ*ibeA*, and **(E)** LF82Δ*ibeA:ibeA* at 2 **(i)**, 24 **(ii)**, and 48 **(iii)** hpi. Representative images for n= 3. Scale bars = 50 μm **(F)** Quantification of neutrophils per intestine for DSS-treated fish infected with above mentioned LF82 strains or paramecia only control. Kruskal-Wallis test. n ≥11. *, P ≤ 0.05; **, P ≤ 0.001; ****P ≤ 0.0001; ns, not significant.

## DISCUSSION

In this study, we present the larval zebrafish as a model to study the interplay between host inflammatory responses and AIEC colonization. During the initial hours post infection, AIEC is observed colonizing the foregut and the midgut, however over the course of infection, AIEC shows a preference for colonizing the midgut of larvae, similar to EHEC (22). This region of the intestine contains absorptive enterocytes, mucin secreting goblet cells, and M-like cells, all of which are also found in the mammalian small intestine (49). Accordingly, AIEC predominantly colonizes the mammalian small intestine compared to the colon of IBD patients (3, 50-52).

By combining a previously published DSS colitis model (17) with food-borne AIEC infection in larval zebrafish, we were able to analyze host-microbe interactions in a dynamic fashion, using intravital and high-resolution imaging of live and euthanized larvae. We observed that AIEC LF82 colonizes better than non-pathogenic *E. coli* in hosts with and without pre-existing intestinal inflammation, which is in accordance with observations reported in murine studies (38). To date it is still unknown whether the colonization of AIEC in a susceptible host triggers the onset of intestinal inflammation or if inflammation presents a favorable environment for the AIEC pathotype. Our data suggest that AIEC persist and promote inflammation in healthy hosts, but is particularly adapted to colonize and persist in hosts with ongoing colitis. The data also suggests that, while uninfected hosts can recover from colitis after removal of pro-inflammatory stimuli (here, DSS), tissue repair and healing as impaired in hosts colonized with AIEC.

There are a few differences between rodent and zebrafish models of AIEC. Although both mice and larval zebrafish have an endogenous microbiota, the rodent microbiome renders mice highly colonization resistant, and AIEC models involve antibiotic treatment to remove much of the endogenous microbiome to allow for AIEC colonization. In contrast, larval zebrafish do not need to be treated with antibiotics to remove the endogenous microbiome, and a single dose of 10^4^-10^5^ CFUs of AIEC consumed through food-borne infection is sufficient to promote bacterial colonization. Mice are usually challenged with 10^8^-10^9^ CFUs of AIEC through oral gavage daily for 3 or 15 days, making them more labor intensive (8, 39, 53). The existing zebrafish model of AIEC infection only requires bath immersion, but adult zebrafish are required to achieve robust colonization (16). Here, we found that AIEC colonization causes increased mortality in DSS colitis fish, compared to unfed or paramecia fed DSS treated fish (**Fig. 5H**). This is consistent with mouse studies, where AIEC LF82, but not *E. coli* strain K 12, decreases the survival of CEBAC10 and DSS-treated beginning at 2 dpi, and by 7 dpi the survival rate of the host is 20% (8).

We found that AIEC LF82 exacerbates intestinal inflammation in hosts with pre-existing inflammation. This is supported by an increase in neutrophil recruitment to the intestine, the inability of the mid-intestine to heal while colonized with AIEC, and the increased relative expression of the genes encoding the pro-inflammatory markers *cxcl8, il1b*, and *mmp9*. Cxcl8 is primarily associated with the activation and mobilization of neutrophils, whereas Tnfα and Il-1β are involved in signaling pathways that regulate apoptosis and cell survival (54). Mmp9 degrades the extracellular matrix during inflammation and through this process activates cytokines that mediate tissue and wound healing (55), however its activation can also contribute to intestinal damage during IBD (56).

To investigate whether *fimH* and *ibeA* are important for the colonization of AIEC in the zebrafish intestine, these 2 genes were deleted from the parent strain. These two genes have been previously characterized *in vivo* and *in vitro*, and thus we reasoned that characterizing their phenotypes would allow us to benchmark our model against published *in vivo* and *in vitro* AIEC models. Prior studies show that the burden of AIEC LF82Δ*fimH* is significantly decreased at 2 and 10 dpi in two different mouse models that express mammalian CEACAM6 in the intestine (8, 57). Deletion of *ibeA* did not impact the burden of AIEC strain NRG857c in mice, although it did contribute to invasion and intracellular survival *in vitro* (20). In the larval zebrafish model, deletion of either *fimH* or *ibeA* transiently caused a higher bacterial burden early during infection, but decreased bacterial burden at later time points (Fig. 6, 7). A possible reason is that LF82 express additional virulence factors involved in adhesion, including OmpA, OmpC, long polar fimbriae, and the lipoprotein NlpI (58-62). Alternatively, the transient increase in burden could be due to an altered immune response, since both FimH and IbeA are involved in neutrophil recruitment and pro-inflammatory signaling in our model (**Fig. 8**). It is possible that *fimH* or *ibeA* deletion cause a defect in bacterial clearance early during infection, and adhesion and invasion defects during later time points. In addition to their immunogenicity, FimH and IbeA both played a role in sustaining epithelial damage and prevention of healing in DSS colitis fish. It is likely that their role in pro-inflammatory signaling and in blocking tissue recovery are linked. Our findings are in line with other studies that show that the colonic epithelium of mice infected with LF82Δ*fimH* and NRG857cΔ*ibeA* appears less corroded than that of mice infected with the parent strains (8, 20).

Recently published work established adult zebrafish as a model of AIEC infection and showed that adult zebrafish produce S100A-10b, a protein homologous to calprotectin, in response to intestinal inflammation caused by LF82 (16). This recent study further supports the observation that AIEC induce inflammation in zebrafish. The decision to use adult or larval zebrafish to study AIEC depends on the type of readouts required to address a question of interest. In contrast to larvae, adult zebrafish are not transparent, which hinders dynamic imaging of single cells. However, in contrast to larvae adult fish have a functional adaptive immune system, which allows studies on this aspect of host-microbe interactions.

The reason why AIEC colonizes hosts with pre-existing inflammation more efficiently than untreated fish is not well understood, but there are several potential explanations for this observation. First, DSS damages the intestinal barrier and facilitates the adhesion and invasion of AIEC, which results in bacterial localization closer to the epithelial basement membrane (**Fig. 4 Dvi**). As a result, the bacteria are farther away from the lumen and fail to be cleared out by peristaltic contractions (63). Within the basement membrane, fibronectin, collagen types IV, VII and XVIII, and laminin are abundant, and these host proteins are all known to bind several bacterial adhesins (64). A second reason may be that DSS changes the composition of the intestinal microbiota that may otherwise limit AIEC colonization. Studies show that the administration of the colitis inducing drug 2,4,6-trinitro-benzene sulfonic acid to larval zebrafish changes the proportion of species belonging to the Proteobacteria and Firmicutes phyla (65). Third, intestinal inflammation may cause the overexpression of a receptor important for binding of AIEC. *In vitro* studies suggest that AIEC can increase the expression of host adhesin receptors. For example, the binding of LF82 through FimH to CEACAM6 induces blebbing of apoptotic cell-derived membranous vesicles, which exposes oligomannosidic glycans that serve as AIEC binding sites (66). Moreover, the expression of CEACAM6 is increased by TNFα and IFN-γ following AIEC infections (67). CECAM receptors are cell-surface glycoproteins expressed in epithelial, endothelial, and myeloid cells (8) (68). To date, twelve human CEACAM receptors have been identified and fully characterized (68). In contrast, ten putative CEACAM receptors have been identified in the zebrafish genome and only one CEACAM protein (CEACAMz1) has been characterized. CEACAMz1 is predominantly expressed in gills and to a lesser extent in the intestine (69). Interestingly, mammalian CEACAM6 is also expressed in the alveolar and airway epithelial cells of the lungs under homeostatic conditions and is highly expressed in the gut only during intestinal disease (70). Furthermore, larval zebrafish express a CEACAM6-like protein (encoded by the *zgc:198329* gene) in the intestine that is 29% identical to human CEACAM6 (71). Future studies are required to investigate to what extent CEACAM proteins are involved in the binding of AIEC in the zebrafish intestine.

We propose that this model may be used identify a common AIEC molecular genetic signature in genotypically diverse strains and to provide a means to develop diagnostics and alternative therapeutics for IBD patients. It has been argued that a plausible reason that such molecular markers have not yet been discovered arises from the limitations of currently used infection models and *in vitro* models to classify *E. coli* strains as AIEC (72, 73). Previous attempts to identify molecular markers of AIEC have relied on *in vitro* systems to quantify adhesion, invasion, and replication inside of infected cells, since there are no widely conserved genetic features, such as the LEE pathogenicity islands in EHEC/EPEC, or certain toxins, such as in the case of STEC (Shiga-like toxins) and ETEC (LT/ST enterotoxins). However, it is plausible that there may be genes essential for AIEC *in vivo* colonization that are not expressed in a simplified *in vitro* model, or are disproportionately important in facilitating colonization only in hosts with pre-existing inflammation. Comparative transcriptomic studies show that the pathogenicity of AIEC changes when AIEC cells are grown *in vitro* and in the presence of host factors (50, 73, 74). Whether or not AIEC contain specific molecular signatures is not currently known but it has been proposed that there are undiscovered AIEC-specific genes that are not commonly found in non-pathogenic *E. coli* strains that are yet to be identified (75). These are hypotheses that may be addressed using transposon mutagenesis and high-throughput assays in larval zebrafish. We also propose that larval zebrafish may facilitate the screening of drugs that target AIEC. Positive results regarding microbial virulence factors, host factors contributing to disease progression, and initial drug candidates in larval zebrafish, may then be further evaluated in mammals. We expect this to present a cost-effective way to identify novel genes that link AIEC with the development or progression of IBD.

## MATERIALS AND METHODS

### Ethics Statement

Zebrafish care, breeding, and experiments described here are in accordance with the Guide for the Care and Use of Laboratory Animals have been approved by the Institutional Animal Welfare Committee of the University of Texas Health Science Center, Houston, and protocol number AWC-22-0088.

### Zebrafish maintenance and breeding

The zebrafish lines used in this study were wild-type (AB) and transgenic lines Tg(*mpo*::*egfp*) (76) and Tg(*mpeg1*:*egfp*) (77) which express EGFP in neutrophils and macrophages, respectively. Adult fish were kept in a recirculating tank system at the University of Texas Health Science Center at Houston Laboratory Animal Medicine and Care on a 14:10 h light: dark cycle at pH 7.5 and 28 °C. Eggs were obtained from natural spawning of adult fish. Fertilized embryos were bleached for 30 sec. in 0.05% sodium hypochlorite solution (stock 4.00-4.99%, Sigma-Aldrich) and kept at 30 °C on a 14:10 h light:dark cycle at pH 7.4. Embryos were raised in petri dishes containing E3 buffer (10 mM HEPES, 5 mM NaCl, 0.17 mM KCl, 0.4 mM CaCl_2_, 0.67 mM MgSO_4_, pH 7.4). The 1X E3 medium was prepared with 10mM HEPES to neutralize the acidic (pH 3) solution that arose after dissolving DSS in standard E3 buffer. Larvae that were maintained past 6 days post fertilization (dpf) were fed GEMMA Micro 75 (Skretting) until euthanized. The larvae were maintained in in 150 mm diameter petri dish containing 90 mL of E3 medium.

### Bacterial strains and growth conditions

The bacterial strains and plasmids used in this study are listed in Table 1. All strains were grown at 37°C in Luria-Bertani (LB) broth or on LB agar plates, with ampicillin (200 µg/ml), kanamycin (50 µg/ml), chloramphenicol (35 µg/ml), tetracycline (10 µg/ml), or gentamycin (15 µg/ml), when required.

**Table 1.** Bacterial strains and plasmids.

| Bacterial Strains | Relevant Characteristic(s) | Sources/References |
| --- | --- | --- |
| *E. coli* MG1655 | Non-pathogenic lab *E. coli* |  |
| AIEC LF82 | Adherent-invasive *E. coli*, parent strain | Torres lab, UTMB |
| LF82Δ*fimH* | LF82 derivative, *fimH* deletion | This study |
| LF82Δ*ibeA* | LF82 derivative, *ibeA* deletion | This study |
| LF82Δ*fimH*:*fimH* | LF82 derivative, *fimH* complementation | This study |
| LF82Δ*ibeA*:*ibeA* | LF82 derivative, *ibeA* complementation | This study |
| *E. coli* DH5α | Used for cloning experiments |  |
| *E. coli* DH5α λpir | Used for complementation | (79) |
| **Plasmids** |  |  |
| pDOC-C | Cloning vector | (78) |
| pDOC-K | Carries the kanamycin cassette |  |
| pACBSCE | recombineering plasmid, encodes the I-SceI and the λ-Red proteins |  |
| pME6032:mcherry | Encodes mCherry protein |  |
| pSTNSK-Cm | Tn7 transposase expression vector | (79) |
| pGpTn7 | Cloning vector |  |

The LF82 deletion strains were generated using by recombineering, as previously described (78). Briefly, constructs were generated by amplifying a kanamycin cassette from the plasmid pDOC-K using oligonucleotide pairs that contain at least 45 bp of homology to the DNA immediately upstream and downstream of the target genes (Table 2). The amplified fragment was inserted into the plasmid pDOC-C, and the construct sequence was verified by sequencing (Azenta Life Sciences). The constructed pDOC-C deletion plasmid and the recombineering plasmid pACBSCE were co-transformed into LF82 via electroporation and plated on LB agar containing chloramphenicol, ampicillin, and kanamycin. Selected colonies were grown in LB broth containing 0.5% glucose for 2 h and then induced with 0.5% arabinose for 4 h. The cells were then collected by centrifugation, and spotted on LB plates without NaCl, but containing 5% sucrose and kanamycin. Sucrose insensitive and kanamycin resistant recombinant colonies were transferred to LB chloramphenicol plates to confirm loss of the pACBSCE plasmid. Loss of the pDOC-C plasmid was confirmed with pDOC-specific oligonucleotides. Gene deletion was assessed by PCR using primers listed in Table 3.

**Table 2.**
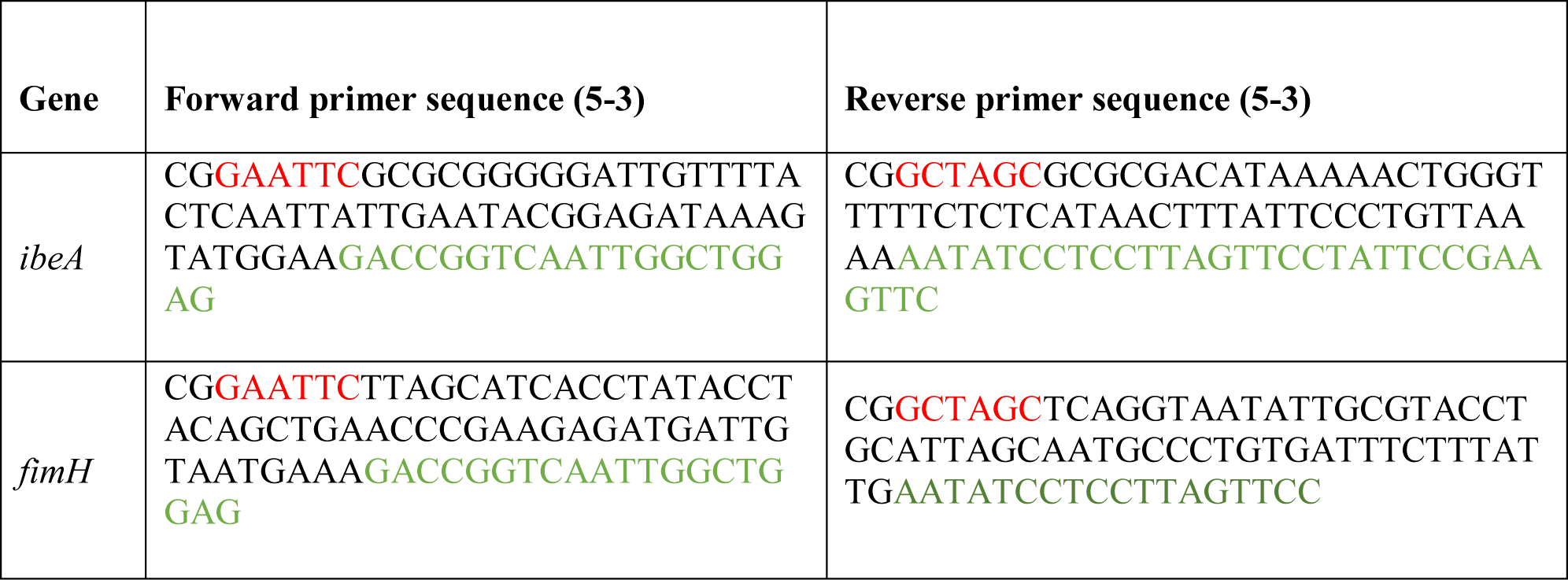
List of primers used to amplify the pDOC-K plasmid with 45 base pair homology (bold) to the DNA upstream and downstream of *ibeA* and *fimH*. The restriction site is in red and the region homologous to the kanamycin cassette is in green.

**Table 3.**
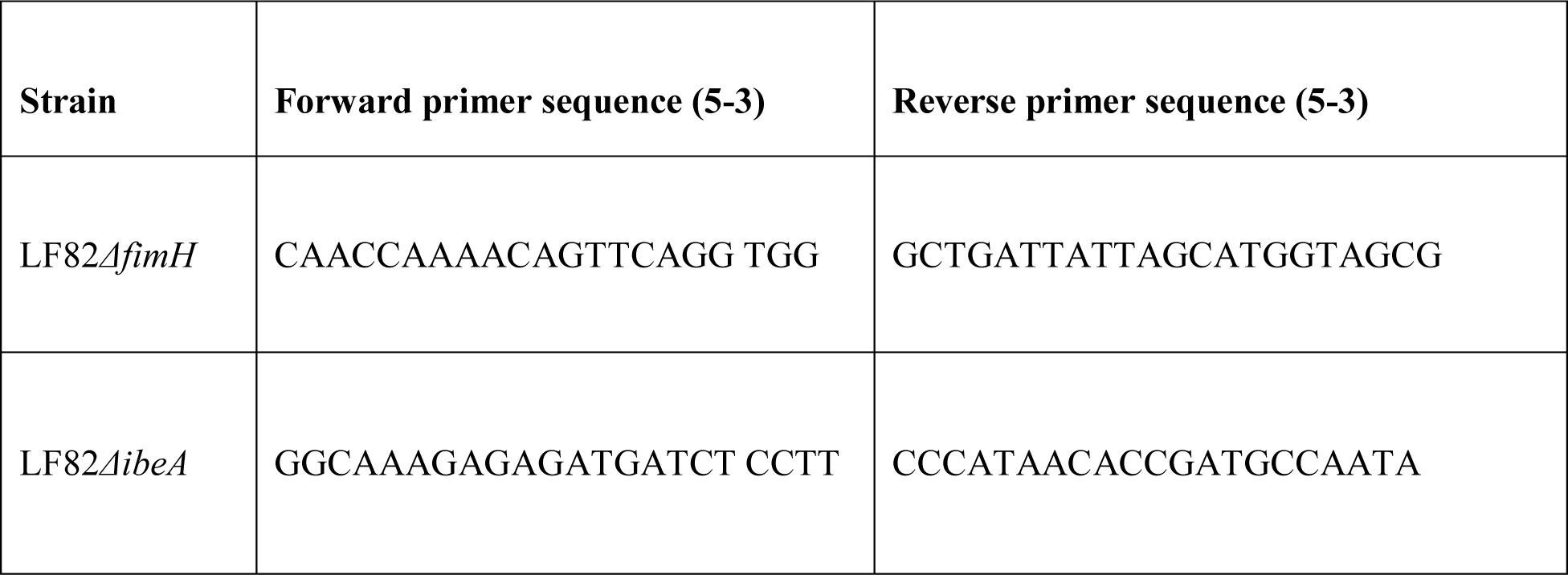
List of primers used to verify deletion mutants.

The complementation strains were constructed by insertion of the gene of interest and its endogenous promoter into the respective deletion strains using a Tn7 based vector system (79). Briefly, the genes were cloned in pGp-Tn7-Gm, and then introduced in DH5α-λpir strain by electroporation to construct pGp-Tn7-fimH and pGp-Tn7-ibeA vectors. Positive clones were checked by colony PCR and confirmed by Sanger sequencing. The pGp-Tn7-fimH and pGp-Tn7-ibeA vectors were electroporated into LF82Δ*fimH* and LF82Δ*ibeA* harboring the Tn7-transposase encoding, temperature-sensitive plasmid pSTNSK-Cm. The cells were spread on LB plates containing gentamycin and chloramphenicol, and then incubated at 30 °C for 20 hours. Selected colonies were further streaked on LB agar plates without antibiotics, and incubated at 42 °C for 20 hours to promote the loss of plasmid pSTNSK-Cm. The colonies were passaged 4-5 times on LB agar plates (no antibiotic), incubated at 37 °C, and screened for resistance to gentamycin and sensitivity to chloramphenicol.

The deletion of *ibeA* or *fimH* and their integration at the attTn7 site was confirmed by PCR (Table 4) and whole genome sequencing. Genomic DNA was isolated using DNeasy Blood and Tissue kit (QIAGEN, catalog no. 69504) and analyzed by Nanopore sequencing (Plasmidsaurus). Plasmidsaurus also generated a complete genome assembly and annotation. Inspection of those genome assemblies showed that the intended mutations were present in the appropriate strains and that the complementation constructs were correctly integrated at the expected loci. To rule out the possibility that fortuitous mutations were introduced during strain construction, two bioinformatic approaches were used. First, we used Snippy (https://github.com/tseemann/snippy) to compare the nanopore reads to the reference genome (composed of the chromosome https://www.ncbi.nlm.nih.gov/datasets/genome/GCF_021398935.1/ and plasmid https://www.ncbi.nlm.nih.gov/nuccore/NC_011917.1/). Second, we mapped the nanopore reads to the same reference genome using Minimap2, and then used FreeBayes to identify possible single nucleotide polymorphisms (SNPs). The candidate SNPs identified by either approach were analyzed by inspecting the alignments with IGV (https://software.broadinstitute.org/software/igv/download). This showed that there were no fortuitous mutations that were introduced during strain construction. The *E. coli* strains were electroporated with the mCherry-expressing pME6032 plasmid, to visualize the bacteria inside of the zebrafish intestine.

**Table 4.**
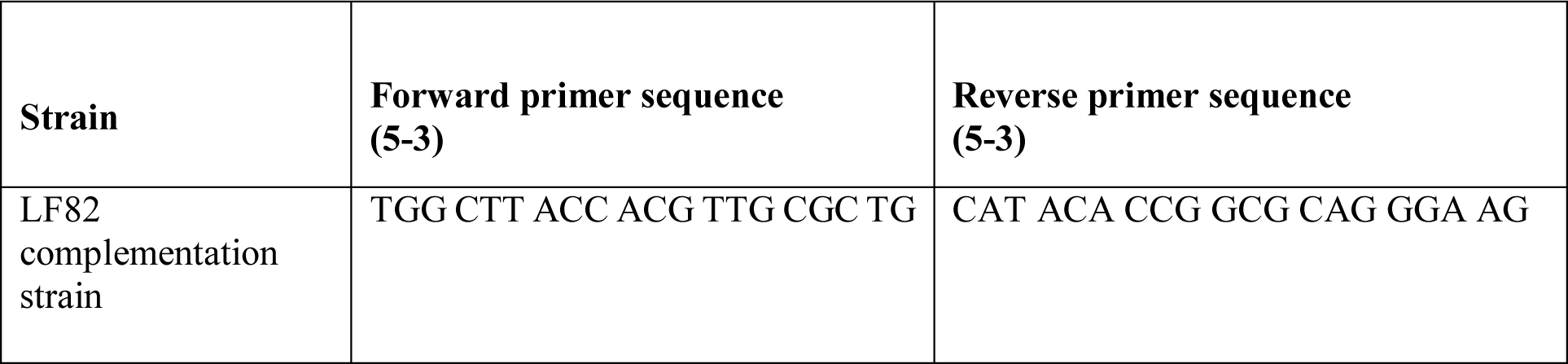
List of primers used to analyze the integration of the Tn7 transposon system at the attTn7 site located downstream of the *glmS* gene.

**Table 5.**
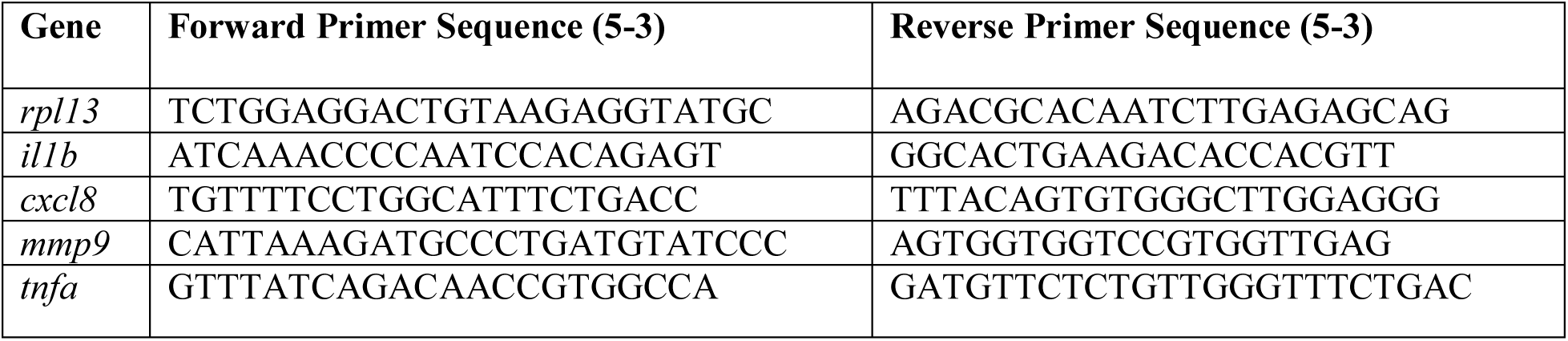
List of primers used to analyze the transcription of pro-inflammatory genes and housekeeping genes.

### Burden of *E. coli* inside of paramecia and larval zebrafish infections

Paramecia were propagated one day prior to the infection experiment and every 2 weeks to maintain live cultures. Loading of paramecia with AIEC LF82 and MG1655 was conducted as described previously (18). On the day of the experiment, paramecia were co-cultured with either AIEC LF82 or MG1655, and the amount of *E. coli* inside of the paramecia was assessed by lysing the paramecia with 1% Triton X-100 followed by colony forming unit (CFU) dilutions and plating, as previously described (18).

The number of *E. coli*-loaded paramecia were counted using an automated cell counter (Life Technologies Countess II) and a final concentration of 2*10^5^ paramecia/mL in E3 medium was used to feed *E. coli* to the larvae for 2 h at 30 °C in a 6-well sterile plate.

### *E. coli* burden and persistence in larvae

The *E. coli* burden in zebrafish larvae was assessed starting two hours post infection (hpi). Briefly, the larvae were anesthetized in the E3 medium with 0.16 mg/mL tricaine, and washed six times to remove excess paramecia. Infected zebrafish larvae were euthanized with 1.6 mg/mL of tricaine. The euthanized larvae were then incubated with 100 μL of a 1 mg/mL filter-sterilized pronase solution, vortexed, and placed at 37 °C for 6 minutes. The larvae were then homogenized by repeated passage through a 31-gauge needle attached to a 1 mL syringe. In all cases, the samples were serially diluted, and 5 μL of each dilution was plated on CHROMagar™ O157 plates (Drg International Inc). The plates were incubated at 30 °C for 24 h, and then at room temperature for an additional 24 h to permit full growth of colonies. The number of dark steel-blue (AIEC) and mauve (MG1655) colonies were assessed afterwards. Data were analyzed with the GraphPad Prism software, version 9.

### DSS administration and survival analysis of DSS-treated larvae

Colitis grade dextran sulfate sodium (DSS) (36,000-50,000 MW, MP Biomedical) was used to induce enterocolitis as previously described by others (17). At 3 dpf, 120 larvae were anesthetized with 0.16 mg/mL of tricaine and transferred to a 150 mm diameter petri dish containing 90 mL of freshly prepared 0.5% (w/v) DSS dissolved in E3 medium. The DSS treatment was followed for 3 consecutive days. Survival or death was assessed daily by observing the presence or absence of a heartbeat on anaesthetized larvae using an Olympus SZX10 stereomicroscope. Dead larvae were removed, and the survivors were transferred to a new petri dish in DSS containing E3 medium every day following assessment.

### Measurement of intestinal and body length, and swim bladder assessment

All larvae were imaged on an Olympus SZX10 stereomicroscope at 1.6 X magnification. Fish were anesthetized in 0.16 mg/mL tricaine and embedded in 1% low melting agarose (LMA). ImageJ was used for image analysis to assess whole body and intestinal length. The length of the intestine was measured from the beginning of the bulb to the end of the cloaca, and the total body length was determined from the mouth to the tip of the tail. The presence of a swim bladder was visualized under the stereomicroscope on anesthetized larvae. The data were analyzed using GraphPad Prism.

### Histological analysis

Zebrafish larvae were fixed in 4% formaldehyde diluted in PBS and incubated overnight (O/N) at 4 °C. Larvae were processed for histological analyses by the UT-Health Core Histopathology Lab. Briefly, larvae were embedded in paraffin, sectioned along the sagittal plane at 2 μm, and stained with hematoxylin and eosin (H&E). Imaging was performed on an AmScope microscope with a MU1003 camera and the AmScope software version x64, 3.7.11443.20180326.

### Neutrophil and macrophage recruitment

Zebrafish larvae were anesthetized, embedded in 1% LMA in a 6-well glass bottom plate, and imaged on an Olympus Fluoview FV3000 confocal microscope for 3-21 hpi. A Z-stack of 190 images of 2 μm slices was analyzed with Fluoview FV3S-SW. The images were then imported into the Imaris software, version 9.7.2, which was used to quantify the number of intestinal GFP-expressing neutrophils or macrophages over the course of 3 to 21 hpi.

### Immunofluorescence

Larvae were euthanized and placed in a 4% formaldehyde solution O/N at 4°C. Then the larvae were washed twice with 1X PBS, permeabilized in acetone for 15 minutes at -20°C, and incubated in PBDT blocking solution [PBS, 1% BSA, 1% DMSO, and 0.5% Triton-X100] O/N. The larvae were then incubated with anti-α-laminin at a 1:25 dilution (Sigma-Aldrich, L9393) O/N at 4 °C. The following day, the samples were washed and incubated with goat anti-rabbit IgG Alexa Flour 488 using a 1:250 dilution (Thermo Fisher Scientific, A27034) and 1 µM/mL 4′,6-diamidino-2-phenylindole (DAPI) O/N at 4 °C. The samples were then washed for 30 minutes, 3 times with a washing solution (1X PBS, 0.1 % Tween-20, and 0.1 % Triton X-100). Some larvae were stained with phalloidin (300 units/mL) and 1 µM/mL DAPI. Samples were imaged on a confocal microscope (Olympus Fluoview FV3000 confocal microscope at 60 X magnification) and images were transferred to cellSENS version 2.3 for deconvolution with five iterations.

### Quantification of bacteria inside of epithelium

Bacteria inside of the intestinal epithelium were quantified on deconvoluted images taken after immunofluorescence imaging. ImageJ was used to quantify the fluorescent signal of the mCherry channel (representing bacteria) (80). The data were plotted using Graphpad Prism and significance was determined using a Mann-Whitney U test.

### RNA isolation, reverse transcription, and quantitative PCR

RNA was isolated from 15 zebrafish larvae for each condition. Briefly untreated or DSS-treated larvae, fed or unfed paramecia, were euthanized, homogenized in TRIzol reagent (Thermo Fisher, 15596026) using a disposable pellet pestle (Fisher Scientific, 12-141-364) and RNA was extracted using a standard protocol (81). Isolated RNA was treated with RNase-free DNase (Qiagen) and cleaned and concentrated using a Zymo Research RNA clean & Concentrator Kit. Removal of DNA contamination was verified by PCR using purified RNA as template.

Reverse transcription was carried out using oligo (dT) primers and the SuperScript™ IV First-Strand cDNA Synthesis Reaction system. The concentration of the cDNA was measured using a Nanodrop-spectrophotometer and 45 ng of cDNA was used for each reaction. cDNAs and primers (listed in Table 4) were mixed with Luna Universal qPCR Master mix (New England Biolabs) and amplification was carried out in duplicate in a CFX96 Real-Time System C1000 Touch Thermal Cycler (Bio-Rad, Hercules, CA, United States). The *elfα* and *rpl13* genes were used as internal controls, and the relative fold-change for each gene of interest was expressed in 2^-ΔΔCT^, where ΔΔCT = [(CT gene of interest-CT internal control) one condition - (CT gene of interest- CT internal control) another condition (82). For DSS experiments, the DSS data were normalized to the untreated group, whereas in the infection experiments the data were normalized to controls fed paramecia without added bacteria.

## ACKNOWLEDGEMENTS

We thank Alfredo Torres (UTMB) for sharing with us the AIEC LF82 strain and Peter Rady (UTHealth Houston) for microscope use to image the histology slides. We also thank Melissa Stephens and Michelle Nguyen from the UTHealth Histology Core for staining and sectioning the paraffin embedded larvae. This work was supported by National Institutes of Health grant R01AI132354 to AMK, and bioinformatic analysis of whole genome sequencing was supported by National Institutes of Health grant R35GM141710 to AvH.

